# Segregation, connectivity, and gradients of deactivation in neural correlates of evidence in social decision making

**DOI:** 10.1101/2020.02.04.934836

**Authors:** Roberto Viviani, Lisa Dommes, Julia E. Bosch, Karin Labek

## Abstract

Functional imaging studies of sensory decision making have detected a signal associated with evidence for decisions that is consistent with data from single-cell recordings in laboratory animals. However, the generality of this finding and its implications on our understanding of the organization of the fMRI signal are not clear. In the present functional imaging study, we investigated decisions in an elementary social cognition domain to identify the neural correlates of evidence, their segregation, connectivity, and their relationship to task deactivations. Besides providing data in support of an evidence-related signal in a social cognition task, we were interested in embedding these neural correlates in models of supramodal associative cortex placed at the top of a hierarchy of processing areas. Participants were asked to decide which of two depicted individuals was saddest based on information rich in sensory features (facial expressions) or through contextual cues suggesting the mental state of others (stylized drawings of mourning individuals). The signal associated with evidence for the decision was located in two distinct networks differentially recruited depending on the information type. Using the largest peaks of the signal associated with evidence as seeds in a database of connectivity data, these two networks were retrieved. Furthermore, the hubs of these networks were located near or along a ribbon of cortex located between task activations and deactivations between areas affected by perceptual priming and the deactivated areas of the default network system. In associative cortex, these findings suggest gradients of progressive relative deactivation as a possible neural correlate of the cortical organization envisaged by structural models of cortical organization and by predictive coding theories of cortical function.

## Introduction

Several proposals on the organization of brain function have suggested the existence of a late stage in supramodal associative processing characterized by distributed parallel computations, in contrast to the predominantly hierarchical character of early stages. Goldman-Rakic (1988), for example, based on anatomical connectivity data from nonhuman primates, drew a contrast between hierarchical early processing and the long-range connectivity of late associative areas forming distinct, reciprocally connected networks (see also Mesulam 1998). Recently, structural and connectivity data in man have been shown to be consistent with these suggestions (for review, see Huntenburg et al. 2018). To date, however, functional imaging studies have made only limited contact with these models. Completely independently, the study of decision making in laboratory animals has uncovered the distributed nature of the high-level neural signal associated with decisions, leading to the suggestion that decision making may provide a general “window on cognition” (Shadlen and Kiani 2013). Based on the assumption that the high-level signal computed in decision making constitutes the late stages of input processing, we pursued the implications of these views of the organization of the brain in a functional imaging study of an elementary social decision task, verifying the consistency of the data with their broad predictions.

Single cell recording studies of perceptual decision making in laboratory animals have demonstrated that a high-level signal is carried by neurons in several regions of the brain, whose firing rates reflect the evidence for the decision (Gold and Shadlen 2002; Cisek and Kalaska 2010; Shadlen and Kiani 2013; Gold and Heekeren 2014). These firing rates may reflect the computation of a decision variable based on the difference between the sensory evidence for the chosen option and for the alternative (Shadlen and Kiani 2013), i.e. the larger this difference, the higher the firing rates. These studies indicate that the evolution in time of these firing rates represents the gradual accumulation of the evidence for choosing among the options on offer until a decision threshold is reached (Gold and Shadlen 2002). Consistently with these findings, functional imaging studies of decision making in man have retrieved sparse cortical signals from the regression of the signal on the difference between the decision values of the chosen and non-chosen options (Heekeren et al. 2008). It has also been argued that this mechanism may also apply to preference-based choice (Krajbich and Rangel 2011; Cisek 2012; Glimcher 2015), and to choices in social cognition (Shadlen and Kiani 2013), where evidence in this respect remains scant (Ruff and Fehr 2014).

More generally, it has been suggested that the evidence accumulation mechanism reveals that brain function may differ in important respects from conventional views in cognitive science (Cisek and Kalaska 2010; Hunt and Hayden 2017). In its purest version, the conventional view envisages informational flow as a hierarchical sequence of processing steps through specialized units (Fodor 1983), whose activation may be univocally interpreted as the recruitment of an identifiable process, and that terminates with the specification of motor output (see, for example, Petersen et al. 1989). Applied to decision making, the sequence of processing steps would consist of encoding sensory information, categorizing this evidence, computing the decision, and sending the outcome of this computation to the motor module (Figure 1a; for discussion, see Heekeren et al. 2008). An ‘alternative view’ is that the areas in which firing rates are associated with the evidence for a decision are part of a dynamically configured network of connected areas in which the information coming from neurons lower in the processing hierarchy and goal representations are integrated, computing evidence for decisions in parallel through the competition of representations of alternatives (Figure 1b; Heekeren et al. 2008; Cisek 2012).

**Figure 1.**
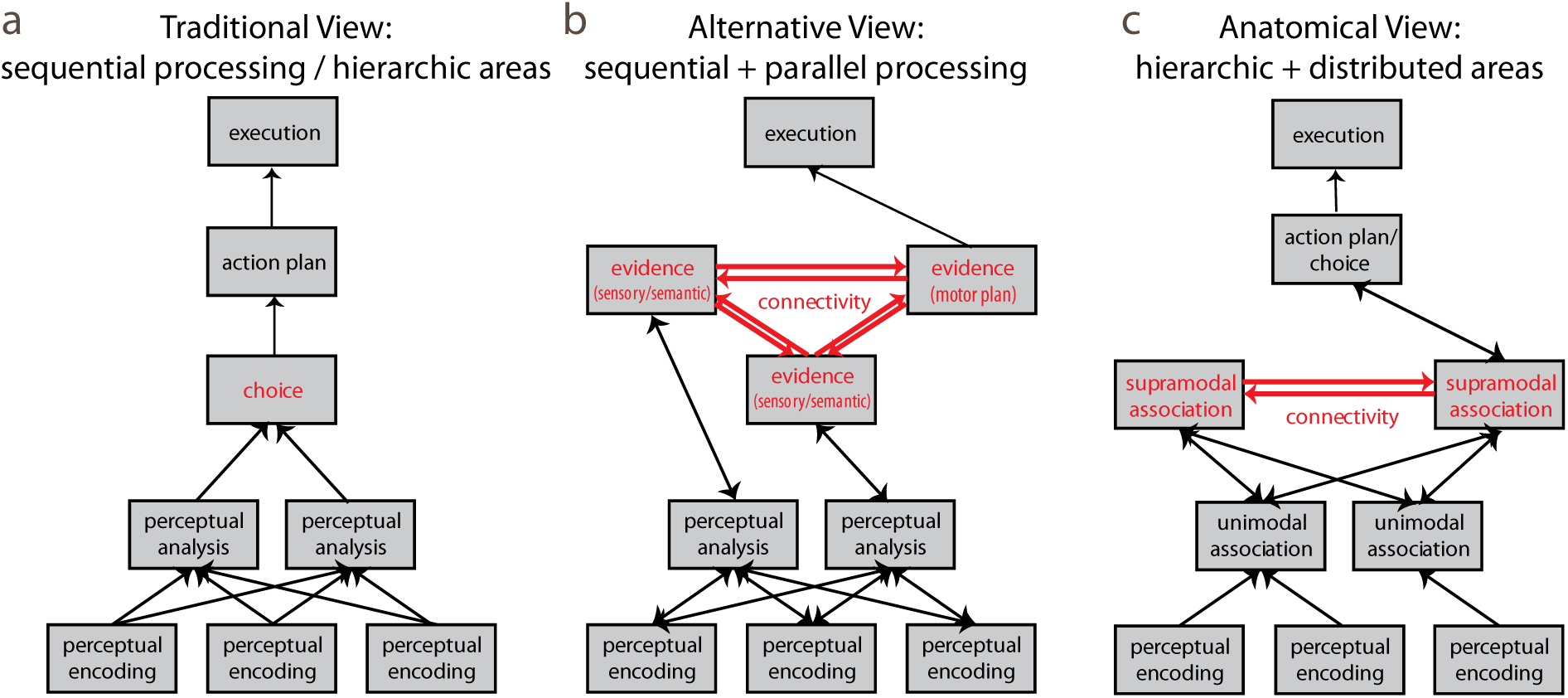
Simplified schemas of three views of cortical organization, highlighting the differences in hierarchical organization and connectivity between the conventional view of information processing and the shared elements of the decision making and anatomical connectivity models. a: in its purest form, the conventional view envisages information processing as a sequence of distinct steps with action planning following input analysis. b: a possible alternative architecture is characterized by accumulation of evidence up to an interconnected network where the decision is elaborated (modified from Cisek 2012). Choice is implemented by a distributed network of ‘evidence hubs’ (in red in panel b) whose activity increases with evidence until a threshold is reached. c: a feedforward information processing model based on anatomical and neuropsychological data (drawn after figures 1 and 3 of Mesulam 1998). Apart from the role of premotor cortex, the decision making and the anatomical connectivity models share the notion of a distinct key role of a distributed network of supramodal areas (in red in panels b and c).

A key aspect of the alternative view is that several areas compute the evidence for the decision simultaneously from multiple modalities. We will refer to these cortical areas as the evidence ‘hubs’ where information relevant for the decision is accumulated in the form of differential evidence for one or the other options. This view integrates segregation (the hubs where evidence for a decision is accumulated are anatomically distinct, presumably reflecting information of different types or modalities) and connectivity (these hubs must talk to each other if all aspects contributing to the decision criterion are to be considered) differently than in the conventional view. In the alternative view, convergence of information in a feedforward cascade of hierarchical steps is complemented by the emergence of distributed computations in connected parallel systems. The discrepancy is best exemplified by motor control. Because accumulation of evidence also takes place in areas deputed to motor control, it is simultaneously available as a motor plan corresponding to the available options.

The present special issue on “gradients in brain organization” is occasioned by recent advances in mapping the large-scale organization of cortical areas within processing hierarchies spanning unimodal sensory regions at one end to transmodal integration regions on the other end (Huntenburg et al. 2018). These gradients become manifest when morphological features of the brain such as cortical thickness (Wagstyl et al. 2015), cortical myelination (Huntenburg et al. 2017, Viviani et al. 2017), cortical myelination and gene expression (Burt et al. 2017), or macroscale connectivity (Margulies et al. 2016) are assessed on a large scale. Comparison with neuropsychological models (Mesulam 1998) supports the correspondence of this gradient with increasingly abstract representations of sensory features (Figure 1c), as well exemplified in the visual and auditory domains (Margulies et al. 2016; Huntenburg et al. 2018). Furthermore, it has been noted that the terminal regions of these gradients overlap with the default network system (Margulies et al. 2016), supporting the suggestion of the involvement of the latter in conceptually-guided cognition (Murphy et al. 2018). Because the default network system is commonly deactivated during the execution of tasks relative to a resting baseline, this observation raises the issue of the relationship between abstract cognitions and deactivations. In the logic of cognitive subtraction used in functional imaging (Posner and Raichle 1994), this would mean that abstract cognition is turned off in all those tasks where these deactivations are seen (i.e. in most tasks, Shulman et al. 1997).

Even if the theoretical premises and the data at the origin of models of decision making and of the organization of supramodal associative cortex differ entirely, there are important elements of convergence in the questions they raise about the large-scale organization of the brain. In the ‘alternative view’ model of decision making of Figure 1b, the cortical hubs where evidence for a decision is accumulated are thought to compute a low-dimensional variable summarizing complex sensory information coming in from a hierarchy of sensory processing areas (Shadlen and Kiani 2013). In the gradient model of brain organization (Margulies et al. 2016) and its precursors (Goldman-Rakic 1988; Mesulam 1998), transmodal association areas are the candidate areas to host this computation (Figure 1c). In the present study, we inquired if the functional imaging of decision making may open a window on the large-scale cortical organization in sensory processing as suggested by the shared elements of the ‘alternative view’ and the cortical gradient models. We first identified the cortical hubs associated with decision evidence in an elementary social decision task, in support of the generality of the evidence signal outside the domains of sensory or preference-based decision making. Subsequently, we attempted to characterize the position of these hubs in the information flow of the model by looking at their relationship with the information they may summarize from areas below, their relationship with task deactivations, and their reciprocal connectivity.

In our experiment, we asked participants to indicate the saddest individual from those shown in two pictures. Participants were divided into two groups, who viewed either facial expressions of varying degrees of sadness or stylized drawings of individuals without facial features depicted in scenes of mourning or in neutral scenes (Labek et al. 2017). When deciding between faces, participants can rely on purely sensory information to make their choice (Vuilleumier et al. 2003). In contrast, the scenes of mourning individuals provided minimal visual detail, and require considering the scenic context or inference about the mental state of the depicted individual to judge between degrees of sadness (Labek et al. 2017).

Our experiment was designed to address three related issues. The first was the detection of the neural correlates of evidence for a decision in an elementary social cognition task. The second was demonstrating segregation and connectivity in the areas associated with the evidence for these decisions. Because the kind of evidence participants based their decisions on differed in these two groups, we expected to detect differences in the hubs between the groups, reflecting different sources of sensory and contextual processing. To assess connectivity, we selected the largest peaks of the evidence hubs associated with accumulation of evidence in associative cortex and used them as seeds in a database of connectivity data (Yarkoni et al. 2011). These seeds were selected in the hubs that differed between input types, so as to ensure that they were related to input processing. Here, we hypothesized that the evidence hubs would be selectively linked by strong connectivity to support integration of the information for the decision, as shown in the networks of the ‘alternative view’ model of Figure 1b.

The third issue concerned the location of evidence hubs in the context of the global organization of the fMRI signal in association cortex. We mapped the location of the evidence hubs with respect of task activations and deactivations, which in the posterior part of the brain formed a gradient from activated perceptual cortex to the deactivations of the default network system. Areas associated with subjective value in preference-based decision making, such as the ventromedial prefrontal cortex (vmPFC) and the postsplenial cortex, are usually deactivated by the task, a fact that has spurred interest in the role of neural inhibition in the constitution of the decision value-associated fMRI signal (Jocham et al. 2012; Hunt et al. 2015). If preference-based and other types of choice making are related, we would expect this relationship to deactivations to occur in both. Indeed, functional neuroimaging studies of negative emotions and emotion regulation often deliver results in vmPFC and in associative cortical areas that combine correlates of regulation with task deactivations so as to challenge conventional interpretations (Benelli et al. 2012; Viviani 2014; Messina et al. 2016). Moreover, as we have already noted, the gradient model of brain organization postulates an integrative role for the nodes of the default network (Margulies et al. 2016), which are usually deactivated during the execution of tasks relative to the resting baseline.

We found that the evidence hubs in associative cortex were typically located in proximity of task deactivations, at a point along a gradient from activation to deactivation. In the sensory-selective network recruited by deciding about faces, its hubs spanned the transition zone between task activations and deactivations, which formed a gradient of activity across the hubs. To verify that the collocation of these hubs in proximity of deactivations was not incidental, we looked at whether task deactivation in the faces group in correspondence of the face hubs differed from that of the mourning group, where these hubs were absent. This analysis will provide suggestive evidence that these areas are embedded in a global organization of information processing whose functional imaging correlate is a gradient of decreasing activation.

Finally, we turned to the temporal dynamics of the signal and the correlates of repetition suppression to identify areas encoding the items of the decision task. Repetition suppression is the decrease of the signal after repeated presentation of stimuli, a manifestation of the neural adaptation and neural encoding of the shared properties of stimuli (Henson 2003; Grill-Spector et al. 2006; Barron et al. 2016). We hypothesized that the correlates of repetition suppression would be distinct from those associated with decision evidence, but still located on the information processing path from the activated sensory cortex to the partially deactivated evidence hubs. In this analysis, we also discovered that, while the network of faces-specific evidence hubs remained stable during the task, there were evidence hubs recruited by mourning scenes that evolved during time. In the initial phase of the experiment, evidence for deciding between mourning scenes activated temporo-parietal and medial prefrontal areas that coincided with task deactivations.

Taken together, the results of the analytic strategy we followed here raise the possibility that, in so far as an fMRI signal referring to a global organization of information processing may be detected in the data of this study, this signal consisted of a gradient of progressive deactivation that was modulated in turn by the existence of the evidence hubs. A consequence of this would be that lack of activation cannot be equated to lack of recruitment. These findings raise the question of whether relative deactivation has any plausibility at all as a neural correlate of information processing. In the discussion, we will mention some independent proposals on the organization of cortical processing that are relevant to this issue. Cortical inhibitory mechanisms arise naturally in neural models that implement competitive weighting of evidence, or in models of cortical function that focus on prediction as a key computational task. This suggests that the progressive task deactivation reported here, while usually a neglected aspect of neuroimaging analyses, is not a finding at odds with current reasoning about cortical function.

## Methods

### Task and modelling

Two groups of participants were asked to indicate the saddest individual depicted in two separate images. In the first group, participants decided on the basis of two pictures of faces displaying varying degrees of sadness (Viviani et al. 2018). In the second group, participants decided on the basis of stylized drawings of individuals in situations ranging from neutral to extreme sadness (mourning the loss of a loved one, Labek et al. 2017). Using two separate groups of participants ensured that the strategy used in choosing between the alternatives was not biased by the fact that participants may have been exposed to both tasks, realizing the relationship between them. The faces were displayed in frontal view from the Karolinska Directed Emotional Faces inventory (KDEF, Lundqvist et al. 1998), and were chosen so as to display varying degrees of sadness range from the neutral to the desperate. The scenes of mourning individuals and neutral scenes were taken from Labek et al. (2017) (Figure 2).

**Figure 2.**
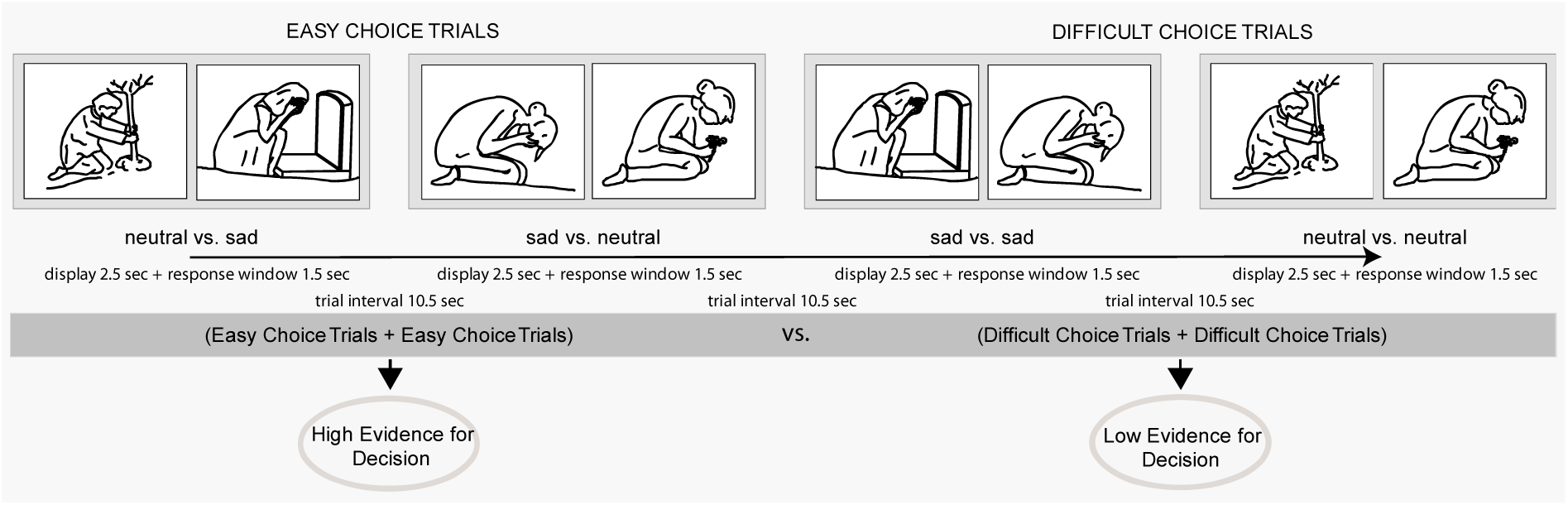
Examples of trials in the mourning pictures group, showing trials where the evidence for the decision is large (easy trials, left) and small (difficult trials, right). Because the same images are presented in all combinations, all images appear in easy and in difficult trials equally. In the faces group (not shown here), the trials were organized in the same way, with faces with different degrees of sadness replacing the mourning pictures.

In each trial, two pictures were displayed side by side on a screen on the back of the scanner visible through mirrors mounted on the coil. All possible combinations of the pictures from a total pool of nine, taken two at a time, were shown (all combinations of the faces pictures in the faces group and all combinations of the mourning scenes pictures in the mourning group). This gave 36 trials, presented in a single run. The two pictures were presented for 2.5 sec, followed by the appearance of two blue circles on the screen below the pictures. During the combined presentation of pictures and circles, participants had 1.5 sec to press one of two buttons to make their decision, after which the trial was declared as a miss. Trials were presented at variable intervals (10.5 sec on average) according to a random exponential schedule bounded at 9 and 12 sec, for a total task duration of 8 min 50 sec. Participants had the opportunity to familiarize with the task in a brief session prior to scanning. The presentation of trials was programmed in standard software (Presentation 14, Neurobehavioral Systems Inc., Albany, CA).

Several strategies have been used in the literature to investigate the neural correlates of evidence. The procedure we adopted here follows the approach of sensory-based decision making in contrasting easy to difficult decisions (Heekeren et al. 2004), but a similar approach may also be found in several neuroeconomic studies of preference-based decisions (Serences 2008; Boorman et al. 2009; FitzGerald et al. 2009; Hunt et al. 2012). From the neuroeconomic studies we borrowed the notion that the possibly subjective degree of sadness participants saw in the pictures could be inferred from their choices. This is because consistency of answers implies the existence of a partial order in the values assigned to the items between one chooses (Samuelson 1938).

In binary choice, the evidence for the decision is simply the difference between the two values of the criterion based on which the choice is made. Hence, to identify the cortical hubs associated with decision evidence in functional neuroimaging data, the signal may be regressed on this difference. After scanning, the score of the times each picture was selected as the saddest of the pair by the participant was computed. The difference and the sum of the sadness scores of the two pictures on display in the trial were computed. The difference was an approximate estimate of the evidence participants had to make their decision. The sum was an estimate of the amount of the properties that determine the decision that was on display. After centering, these participant-specific difference and sum scores were then used as the ‘parametric modulations’ of a regressor sampling the modeling of the BOLD response at each trial. This means that the model included an effect of trials relative to fixation (referred to as the effect of ‘task’ in the results), modeling the effects of perception and encoding of stimuli as well as the generation of responses common to all trials, and the two parametric modulations, modeling the interaction of this effect with the differences between the scores and their sum. Additional parametric modulations were given by a centered score of the number of times the pictures of the trials had been previously presented, and the interaction of the score with the difference and sum of the sadness scores. The parametric modulations captured variance not explained by the main regressor, which modeled the effect of trials. Trials in which participants gave no response (misses) were modelled separately and considered as a nuisance effect.

This modelling strategy was chosen based on its capacity to control for sensory properties of the stimuli associated with the criterion for the choices, thus minimizing confounding of the criterion on the basis of which decisions are made from the effect of evidence. For example, sadness in facial expression may be confounded with salience (or any other property of the visual stimuli that correlates with sad expressions). However, the effect of evidence was not confounded with salience for two reasons. First, in so far as correlated with the degree of sadness, the salience of the images in the trial is summarized by the sum of the scores of these images. The inclusion of the sum of scores in the model therefore adjusts for salience of the trial or any other property that determines choice. Second, when participants make consistent choices, and there are no misses, then the difference and sum of scores regressors are orthogonal. This fact is difficult to prove in general, but may easily be checked directly for a given number of trials (the Matlab code to do this is provided in Viviani et al. 2019). The orthogonality follows from the fact that each picture is presented in association with all other pictures. Two very sad pictures give the same low difference score as two neutral pictures, while the same pictures, in a different combination, give high difference scores. Because the effect of the stimuli themselves is subtracted away in the regression on the score differences that detects the signal associated with evidence for the decision, this regression detects only evidence-related areas (i.e. degree of evidence for sadness), not areas differentially recruited by the sensory encoding of the stimuli.

An alternative modelling strategy consists of obtaining separate coefficients for the regression on the highest and lowest scores of the two items presented in each trial (for example, Boorman et al. 2009) and taking the contrast high score vs. low score to the second level. This modelling strategy differs from the one adopted here in that an additional degree of freedom is spent to model the slopes of the effects of the low and high score items independently. Using this alternative modelling strategy, we obtained results that were qualitatively identical to those reported in the results section. We used here the score difference strategy because of its better control for the confounding of the choice criterion.

As Heekeren et al. (2004) have noted, when the evidence is large the decision is easy (Figure 2). For this reason, in sensory decision making it is common to refer to this regression as to the easy vs. difficult contrast (Heekeren et al. 2004, 2008). It may be noted that the strategy of contrasting easy with difficult decisions in a neuroimaging study is itself non-standard, because difficult decisions are not a control condition for easy decisions. Standard reasoning, based on the cognitive subtraction logic (Posner and Raichle 1994), would suggest that the neural correlates of decision making are those that increase with the difficulty of decisions (as in studies of cognition, Rypma et al. 1999), not those associated with their easiness (Kayser et al. 2010; for a discussion, see Heekeren et al. 2008). In the present work we looked at whether the strategy of contrasting easy vs. difficult decisions revealed information on the large-scale organization of information processing and highlighted the points where the resulting conclusions may depart from those of the conventional approach.

### Recruitment and image acquisition

The study was conducted at the Psychiatry and Psychotherapy Clinic of the University of Ulm, Germany, after approval by the Ethical Review Board. Healthy participants (N = 63) were recruited through local announcements and admitted to the study after verifying that they met the inclusion criteria (lack of medical or psychiatric pathology, lack of metal implants, pacemakers, or extensive tattoos) and had provided written informed consent. To limit the correlation between the difference and sum of scores regressors, we set a threshold of max 4 missed trials to consider the data as valid, leading to the exclusion of 3 participants in each group. The final sample comprised 27 and 30 participants in the faces and mourning groups, respectively.

Data were collected in a Prisma 3T Siemens Scanner using a T2*-sensitive echo-planar imaging sequence (TR/TE: 2460/30 msec, flip angle 82°, image size 64×64×39 obtained from transversal slices of 2.5 mm with a gap of 0.5 mm, giving an isotropic voxel size of 3 mm). A 64-channels head coil was used with foam padding to minimize head motion.

### Data analysis

Data were analyzed with the freely available software SPM12 (www.fil.ion.ucl.ac.uk/spm). After realignment, normalization, smoothing (FWHM 8 mm), and high-pass filtering (512 sec cutoff), trials were modeled by a boxcar function with fixed trial duration 2.5 sec, convolved with a standard haemodynamic function. Realignment parameters were also included in the model as nuisance covariates. The model was estimated separately in each voxel and included a first-order autoregressive term to model autocorrelation in the residuals. Estimates of the contrasts of interest were brought to the second level to account for the random effect of subjects in a factorial design. At the second level, we report significance tests corrected at the peak and cluster level computed by permutation (8000 re-sampling). Peak-level corrections (also known as voxel-level) achieve strong control of false rejection errors. For cluster-level corrections, clusters were defined *a priori* by the uncorrected threshold *p* < 0.001. In both corrections, the testing family was defined by a mask of gray and white matter determined by majority voting on the segmentation computed in each individual separately as part of the normalization procedure.

In the permutation testing for the null conjunction analysis of §3.5, we combined the permutation distributions at voxel and cluster level of the permutations for the two original contrasts, keeping the larger value or the two at each permutation. This provided correction for the joint testing family from both tests. The test statistic was obtained by taking the smallest *t* values of the parametric maps of the original contrasts. To obtain cluster definition thresholds from the smallest *t* values that are comparable to those used in other tests, the cluster definition threshold was set at *p* < 0.002. In this type of conjunction, the null is rejected if there is evidence for both effects obtaining simultaneously (Nichols et al. 2005).

Connectivity data were retrieved from neurosynth.org (Yarkoni et al. 2011) using as seeds the coordinates of the interaction between evidence and group. The data are based on Yeo et al. (2011) and report functional connectivity at rest from a sample of 1000 subjects from about 18.000 cortical loci. The database contains data adjusted for slice timing, realigned, filtered, and denoised to minimize contributions of nonneural origin. For details, see Yeo et al. (2011).

Overlays for Figures 6b and 7 were obtained using the SPM software. All other overlays were obtained with the freely available software MriCroN (Chris Rorden, https://people.cas.sc.edu/rorden/mricron/install.html) from the statistical parametric maps of *t* values. Figures were then annotated with text and legends in Adobe Illustrator. The hatch pattern of Figures 4 and 8a, b was obtained by drawing a grid and using it as a clipping mask on the original overlay from the statistical parametric maps. The signal course of Figures 3 and 8c was obtained by fitting a Fourier series of 5 basis functions for the 6 time-points in the interval of the plot (software package fda, MATLAB version, Ramsay and Silverman 2005) to partial residuals from the fit, with adjustment for the intercept, for the regressor for the misses (if present), and for the movement covariates. The trials were defined by an interval of 15 seconds after each onset. Ninety percent confidence intervals for the fitted curves were computed point-wise, as customary in functional data analysis (Ramsay and Silverman 2005). Histological area classifications were determined with the Anatomical toolbox for SPM (Eickhoff et al. 2005).

**Figure 3.**
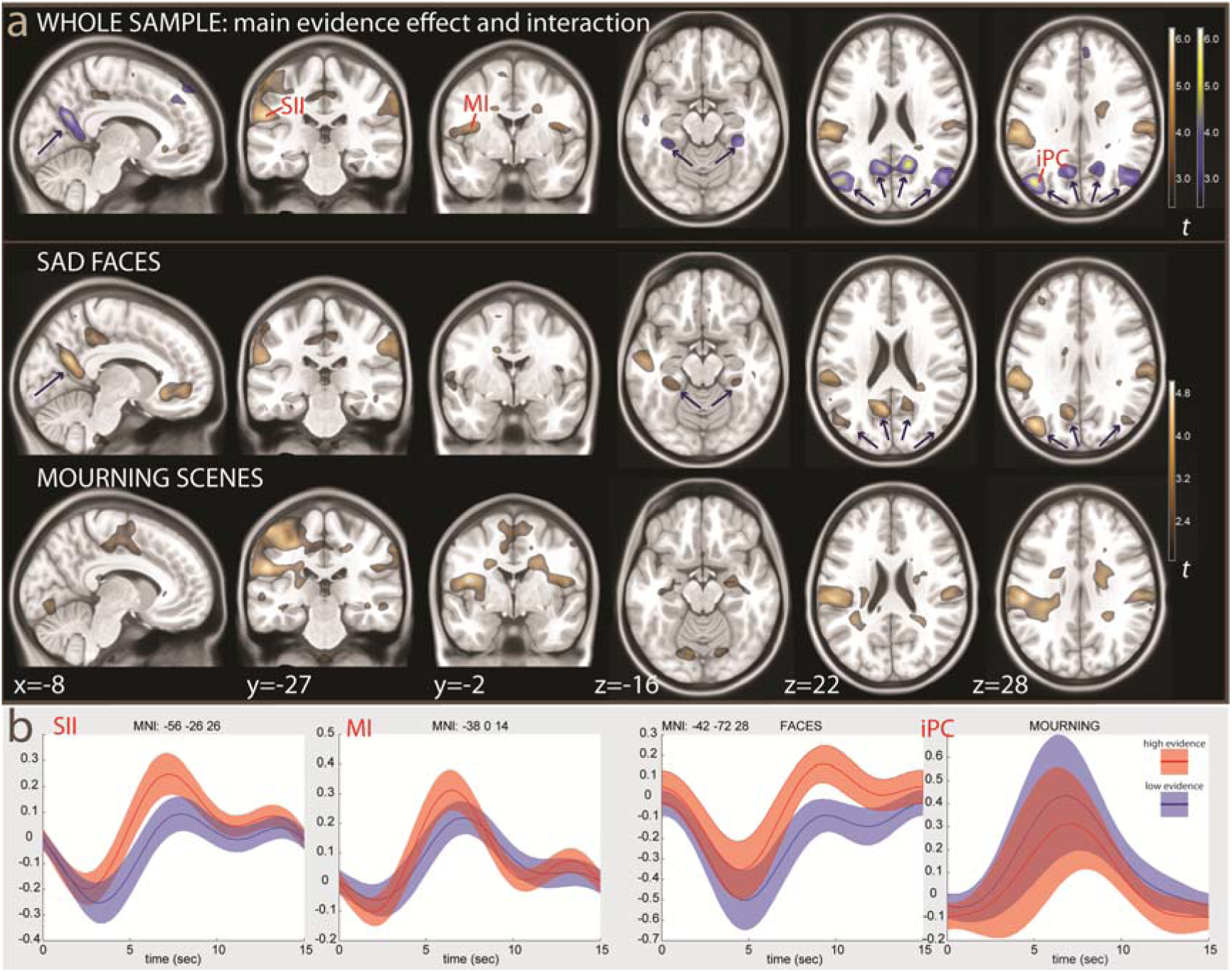
a: the top row shows the main effect of the evidence for decision (in brown) and the interaction with the stimulus type (in blue-yellow). The interactions are highlighted by arrows (overlays thresholded at *p* < 0.01, uncorrected, for illustration purposes). The middle and bottom rows show the effect of evidence in the sad faces and mourning scenes separately (threshold at *p* < 0.05). b: The signal course at the somatosensory and insular peaks (left), and at the iPC/sTG peak (right), where it is shown separately for the faces and the mourning groups. The curves show the signal course and 90% confidence intervals, fitted by a Fourier series, in the tertiles of the trials where the evidence for the decision were highest (easy decisions, red/orange color) and lowest (difficult decisions, light blue color). Coordinates in MNI standard space. SII: sensory association cortex; MI: middle insula; iPC: inferior parietal cortex (posterior portion).

**Figure 4.**
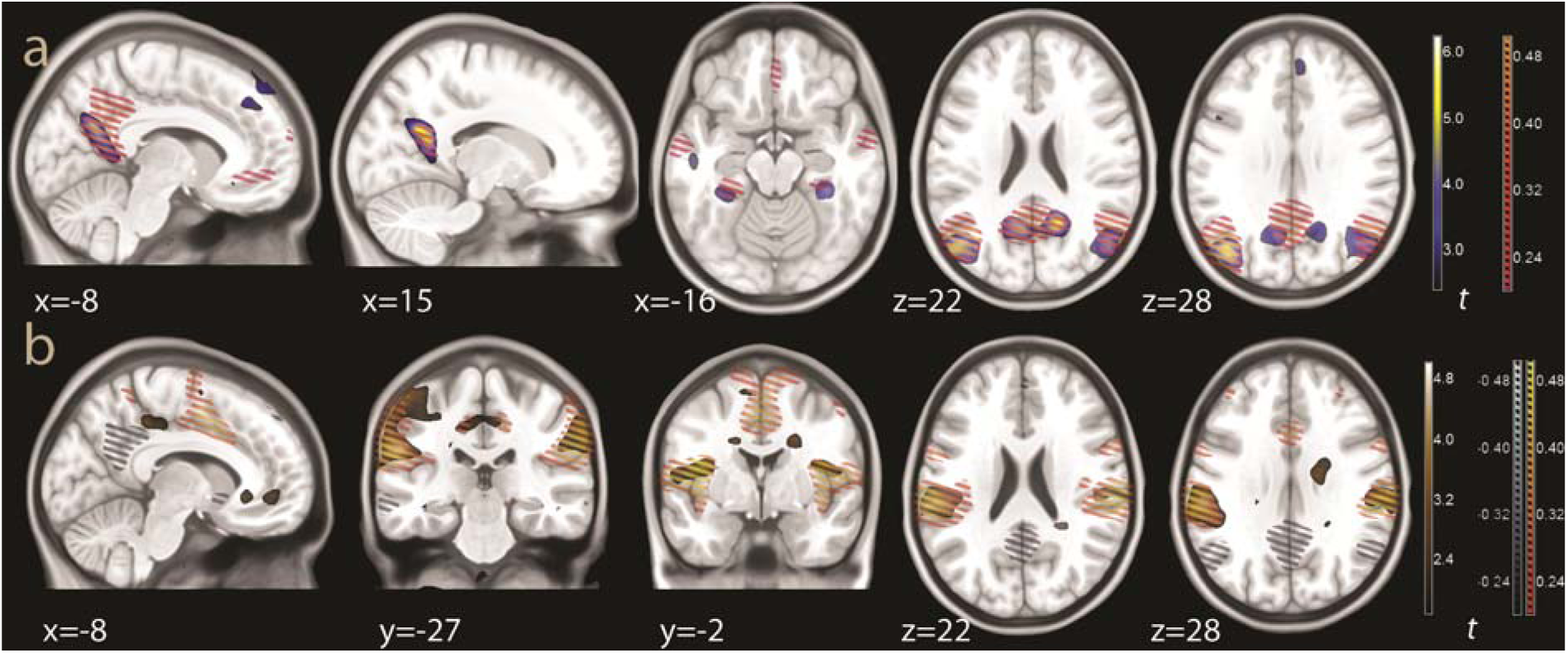
a: connectivity from the left iPC/sTG at −42, −72, 28, overlaid on the effect of evidence for sad faces (interaction evidence × faces). The connectivity is shown as an orange hatch pattern, which largely overlaps with the effect of evidence for sad faces, which is the same effect shown in the top row of Figure 3a (blue to yellow colours). b: connectivity from the left middle insular seed (−26, −16, 16), overlaid on the main effect for evidence (in brown, the same as in the top row of Figure 3a). The positive connectivity is shown as an orange hatch pattern, as in panel a. Negative connectivity is in shades of grey. All connectivity values were thresholded at 0.20, as in the default display from neurosynth.org. Coordinates in MNI standard space.

**Figure 5.**
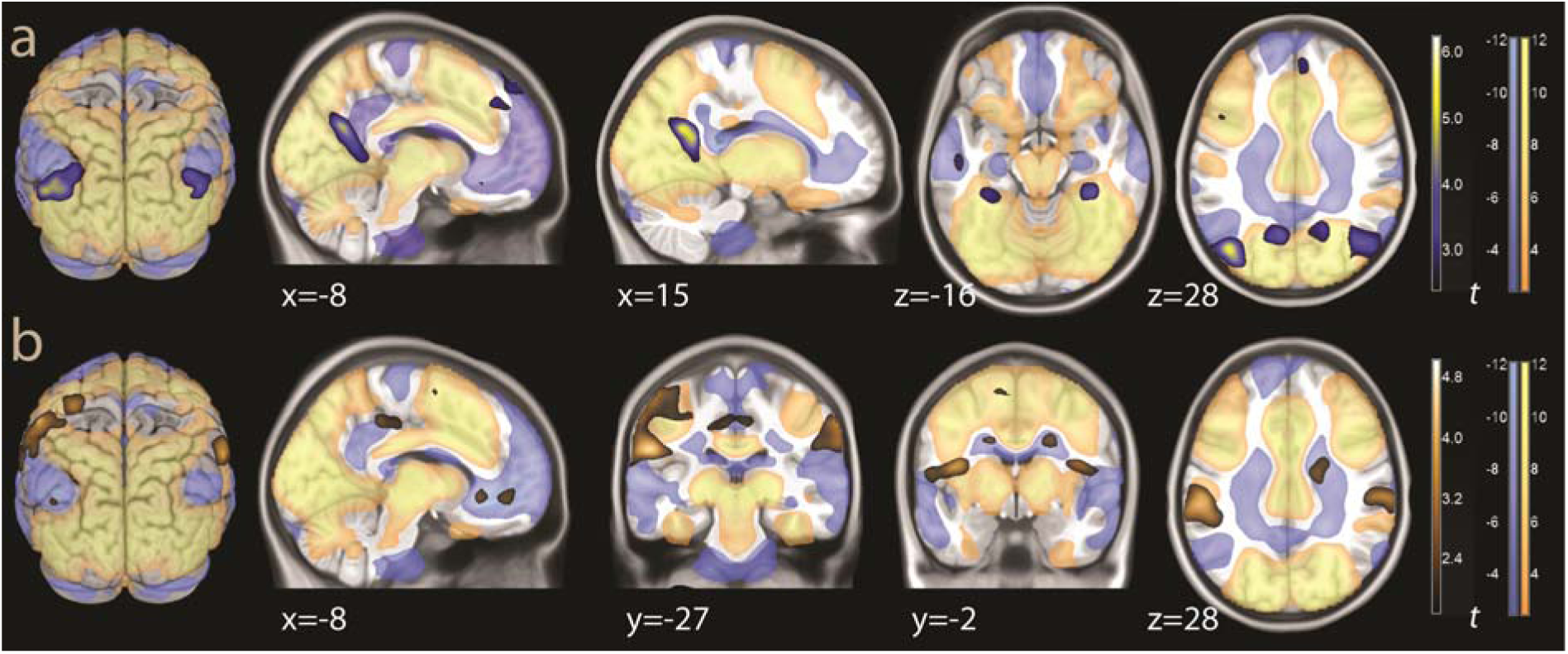
a: overlays for the effect of evidence for sad faces (interaction evidence x faces, same data as in Figure 3, blue-yellow), drawn with overlays of task activation and deactivation (yellow and light blue colors, thresholded for illustration purposes at *p* < 0.001). b: overlays for the common effect of evidence for both groups (same data as in Figure 3, brown colors), together with the same overlays of task activation and deactivation (threshold for illustration purposes *p* < 0.01, uncorrected). Coordinates in MNI standard space.

**Figure 6.**
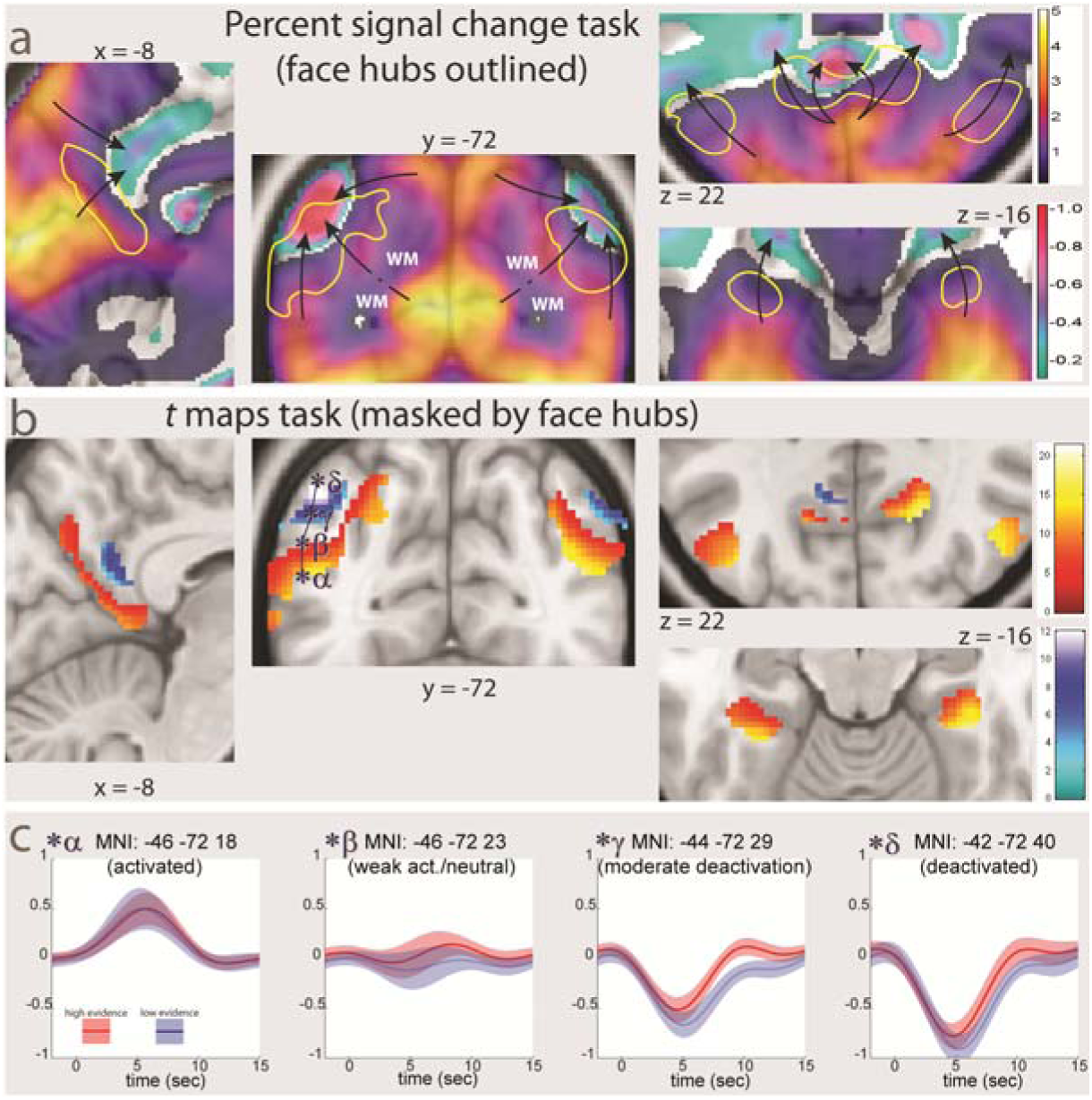
Gradient of progressive task deactivation around and with-in the evidence hubs. a: Percent signal change for the task relative to baseline (thresholded at 0.1%) from sensory to associative cortex. The black arrows show the gradient of progressive deactivation toward the noted of the default network system. In the frontal slice at y = −72, the signal change drops where the slice goes through white matter (WM). The effect of evidence for faces (same data shown in Figure 5a) are outlined in yellow. b: Overlays of *t* maps of task activations and deactivations (in warm and cold colors), thresholded at the uncorrected significance level *p* < 0.001 and masked for the effect of evidence for faces. c: signal course and 90% confidence intervals in the progression from activation to deactivation in the iPC/sTG hub in the faces task (at the points from *α* to *d* marked in panel a, spanning the evidence hub at regular intervals, with *d* at the deactivation peak). In red, signal for trials with high evidence for the decision; in light blue, trials with low evidence. Coordinates in MNI standard space.

**Figure 7.**
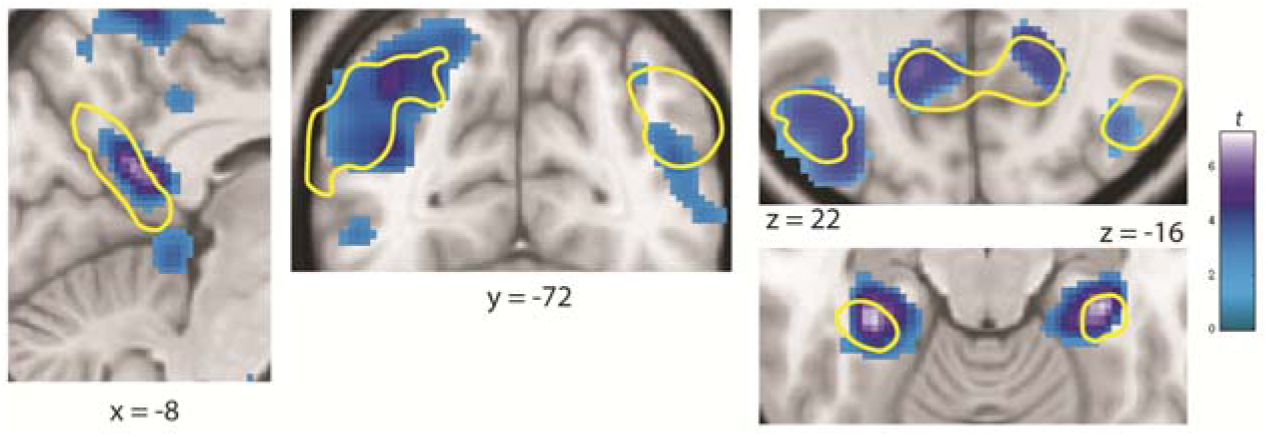
Effect of the contrast faces vs. mourning, showing areas of reduced activity in the faces group thresholded for illustration purposes at p < 0.01, uncorrected, shown in the same locations as in Figure 6. The faces evidence network is out-lined in yellow. Coordinates in MNI standard space.

**Figure 8.**
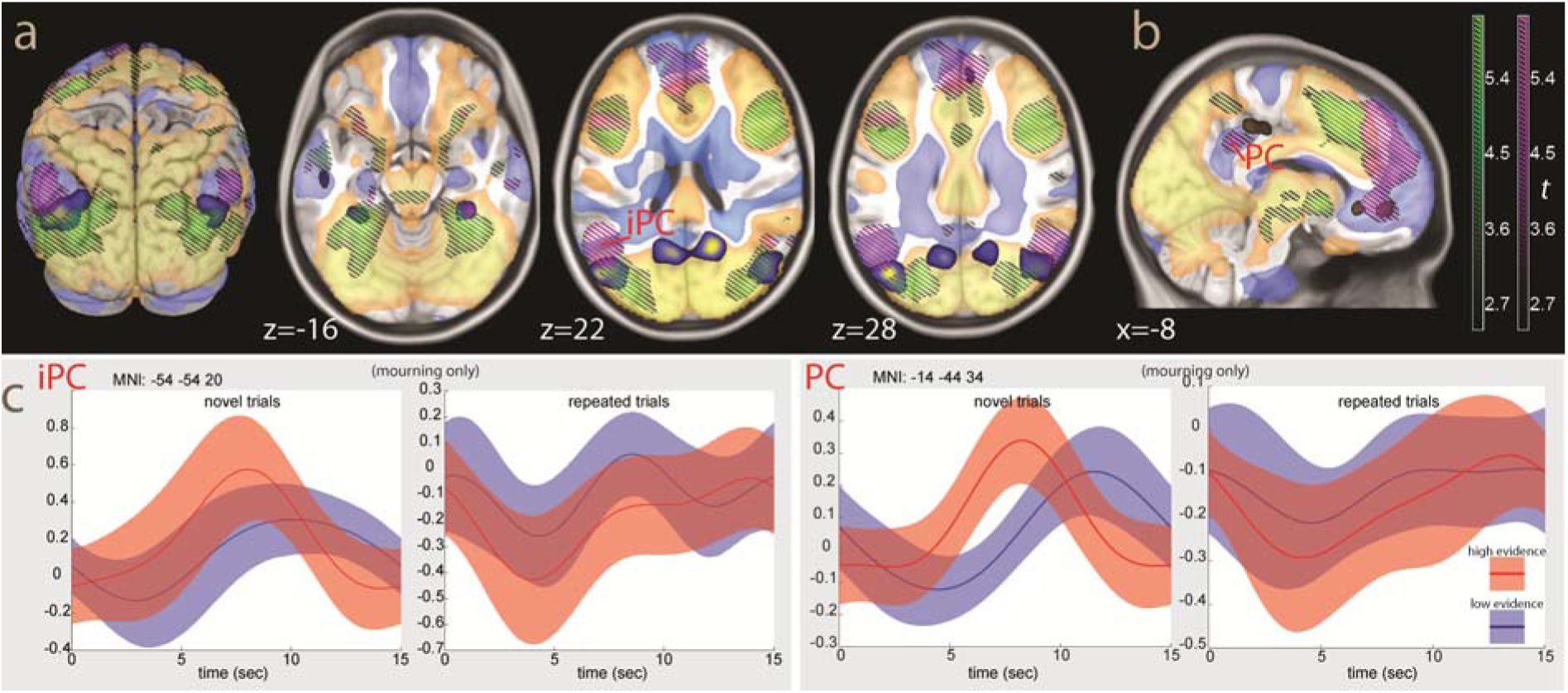
a: overlays for the effect of evidence in sad faces (blue-yellow), drawn with overlays of the effect of repetition (decreases, hatched areas in green) and of the third order interaction evidence × faces × repetition (hatched areas in violet). Task activations and deactivations are in yellow and light blue colors. b: overlays for the common effect of evidence for both groups (brown colors), together with the same other overlays as in panel a (threshold for illustration purposes *p* < 0.01, uncorrected). c: signal course and 90% confidence intervals at the iPS/sTS peak in the tertile of novel and most repeated images trials in the mourning group (left). On the right, the same plots for the posterior cingular locus (PC). The curves show the signal fitted by a Fourier series in the tertiles of the trials where the evidence for the decision were highest (easy decisions, red/orange color) and lowest (difficult decisions, blue color). Coordinates in MNI standard space.

Behavioral data were modelled in the R language for statistical computing (www.r-project.org/) using the *lme4* package (version 1.0, Bates et al. 2015) to model subjects as random effects when appropriate.

## Results

### Behavioural data

Participants gave consistent answers. Emotional items had a higher sadness score (*t* = 15.9, *p* < 0.001), and most subjects chose one picture as the saddest (giving a maximal sadness score of 8), indicating that participants were not choosing the pictures at random. In the sad faces task only two individuals obtained a maximal sadness score of 6, and six a maximal sadness score of 7; in the mourning scenes task, there were one and four individuals with these scores. The difference in the maximal sadness scores in the two groups was not significant (logistic regression with overdispersion, *t* = 0.90, *p* = 0.37). However, there were more misses in the mourning scenes task at trend significance levels (*t* = 2.03, *p* = 0.05), possibly indicating a slightly more difficult task. Response times were negatively associated with the difference in the sadness scores in the trial (*t* = −13.08, *p* < 0.001, repeated measurement regression with subjects as random factor), indicating that responses were quicker in easier trials.

### Evidence for decision (easy vs. difficult decisions)

In the neuroimaging data, we first looked at the effect of the evidence for the decision (easy vs. difficult decision) in both groups taken as a whole, and then at the existence of an interaction between the groups that would support segregation in this system. The evidence for the decision was associated with a signal in a sparse collection of areas that included the sensory association cortex (SII) in the anterior portion of the parietal lobe (areas PFt and PFop in the histological classification of Caspers et al. 2006), the postcentral gyrus (Area 1, Geyer et al. 1999) and the adjacent precentral gyrus, and the middle insula (Figure 3a, in brown colour; see Table A1 in the Appendix).

### Task-specific decision hubs (interaction evidence × group)

In the interaction easy vs. difficult decisions × group, there were bilateral effects in the most posterior portion of inferior parietal cortex/superior temporal gyrus (iPC/sTG), also encroaching into the occipital lobe (area PGp in Caspers et al. 2006) and in the cuneus-precuneus/retrosplenial cortex (Figure 3a, top row, in blue to yellow colours; Table A2). Inspection of the effect of evidence in the two groups taken separately (Figure 3a, middle rows, in brown) revealed this interaction to arise because these areas were recruited during the sad faces task, but not in the mourning scenes task. For this reason, we will refer to this effect as the effect of evidence in faces. Also of note are bilateral effects of evidence in the faces task in the anterior portion of the fusiform gyrus (visible at z = −16 in Figure 3a, area FG4 according to Lorenz et al. 2017). These loci failed to reach significance in the interaction, but we draw attention to them here because they will reappear later in the analysis (see §3.4). In the other direction, i.e. associative areas that were predominantly associated with evidence in the mourning scenes task, there were no significant effects, as the largest peak in this interaction, i.e. the insular peak visible in Figure 3, failed to survive correction for multiple comparisons (*x, y, z*: −26, −16, 16, *t* = −4.08, n.s.). As shown in Figure 3b in these areas the signal showed a more or less marked brief deactivation at the presentation of the stimuli, followed by stronger activity in the trials where the evidence for the decision was stronger.

### Connectivity database search for evidence hubs for faces and mourning

According to the alternative view, the areas where evidence for a decision is accumulated must be talking to each other to integrate different properties of the options and reach a consistent decision. Likewise, anatomical models include distributed computations in the late stages of stimulus processing (Goldman-Rakic 1988; Mesulam 1998). We therefore looked for data on the connectivity of the evidence in faces network in the public database neurosynth.org (Yarkoni et al. 2011). We looked at the connectivity spanned by the left iPC/sTG (seed −42, −72, 28, Table A2), as shown in Figure 4a. In this and the following figures, we show the same data as in Figure 3a (effect of evidence in faces in blue-yellow, effect for evidence in the whole sample in brown) in combination with other effects. In Figure 4a, one can see that the connectivity of this seed, shown in the hatched areas, closely overlapped with the effect of evidence for faces. This overlap also included the anterior portion of the fusiform gyrus that failed to reach significance in the previous analysis (visible at *z* = −16). The only partial mismatch between this connectivity and the effect of evidence for faces concerned the medial face of the brain shown in Figure 4a, (*x* = −8), where the connectivity extended toward areas that were weakly detected by the effect for evidence for the whole group. There was no substantial negative connectivity in the data from neurosynth.org for this seed.

To look for evidence for connectivity in the region associated with evidence in the mourning group, we examined the data for the left insular region (seed −26, −16, 16). We chose this seed because of its importance in the encoding of pain (Peyron and Fauchon 2019), and because it had the strongest activation in the mourning group in the interaction with the effect of evidence. The results, shown in Figure 4b, show again a substantial overlap with the areas associated with the amount of evidence, again with the possible exception of the medial aspect of the brain. Specifically, this connectivity retrieved the activation in the somatosensory cortex detected in the effect for evidence in the sample.

The data on the connectivity for the insular seed also included areas of negative association. These areas are shown as grey hatched areas in Figure 4b. One can see that, in the posterior portion of the brain, these areas of negative connectivity retrieved the areas associated with evidence for faces. In summary, the connectivity analysis showed existence of connections between the peaks activated in the decision task and reproduced the dissociation between the system activated by the information from faces and a more general system active in both tasks. The data also suggest the existence of a third network centered in the posterior cingulus/retrosplenial cortex and vmPFC. While the effects of evidence obtained in this third network were too weak to survive the correction for multiple comparisons, we draw attention to them here because these regions will resurface in the analysis later (see §3.6, interaction evidence × group × repetition).

### Relationship to task activations and deactivations

To embed the evidence networks in the large-scale organization of cortical processing, we examined the relationship between the evidence hubs and task deactivations. In Figure 5, we drew the same effects of decision evidence of Figure 3 together with the activations and deactivations of the task (relative to the implicit baseline of the intertrial intervals). Figure 5 shows that both evidence networks were located toward the end of the areas of activation, in some cases extending beyond it and encroaching the adjacent deactivated areas (in the inferior parietal region visible at *z* = 28 and *y* = −27 of Figure 5). This was especially the case for the faces network, whose hubs were loci located on the ribbon of the transition between activations and deactivations (Figure 5a). The evidence hubs in the medial aspect of the brain were located entirely in deactivated areas. (Figure 5b, *x* = −8).

Figure 6a shows the gradient of progressive task deactivation (in percent of signal change) in the cortex hosting the evidence for faces network, with the areas for the evidence for faces outline in yellow. Figure 6b shows the contrasts for task activations and deactivations in the same region, masked by the same areas detected for the evidence in faces network.

One can see that the location of the evidence network in the transitional zone between activations and deactivations more clearly, and the existence of a centrifugal gradient relative to a point in the occipital pole. Even in the areas in which the evidence networks were in an area activated by the task, the activation effect was modest relative to the main activations. At the point marked by α in the figure, which was activated by the task, the estimated signal change due to the task was 1.48% (standard error 0.14). In comparison, in the left occipital/calcarine cortex ranged between 4.5% and 6% (standard error 0.2-0.3).

Figure 6c shows the progressive modulation of the task vs. fixation signal through the progression from activated to deactivated cortex in and around the iPC/sTG hub. One can see a signal associated with high evidence for the decision where the task activation is weak or there is deactivation relative to the intertrial baseline (loci *β* and *γ* in the Figure). The locus *δ* corresponds to the peak deactivation in the posterior iPC. Here, the signal associated with the evidence for faces is much weaker, if present at all (*t* = 2.77, n.s. after correction).

The location of the evidence hubs at the transition to task deactivations raises the issue of whether the type of input (faces or mourning) changed the level of activity in the task vs. baseline contrast itself. If the evidence hubs are related to a decrement of activation, one would expect activation to be lower in the faces group, as these hubs are present only in this group. This is also the contrast we would adopt in a conventional analysis (even if using faces and mourning scenes as control for each other would not be entirely appropriate), except that we would look for a neural correlate in the form of higher instead of lower activation. We found that in the hubs and the adjacent cortex, activity was lower in the faces than in the mourning group, the opposite of what one would expect if activation in these areas reflected recruitment for the purposes of encoding sensory features (Figure 7; iPC/sTG hub, *x, y, z*: −34, −76, 42, *t* = −4.97, *p* = 0.04; −36, −78, 16, *t* = −5.25, *p* = 0.015; retrosplenial cortex, −14, −56, 18, *t* = −7.10, p < 0.001; 16, −50, 20, *t* = −6.20, *p* = 0.001, all peak-level corrected). In the faces group, activation was lower than in the mourning group also in the anterior fusiform gyrus hub (−30, −36, −16, *t* = −6.75, *p* < 0.001; 32, −32, −14, *p* = −7.21, *p* < 0.001, peak-level corrected).

As Figure 7 shows, there is considerable correspondence between the face hubs (outlined in yellow) and the reduced activity in the faces group. To formally evaluate the co-localization of the faces evidence network and the reduced activity in the faces group of Figure 7, we conducted a null conjunction analysis of these two contrasts (Table A3). After correction, the left iPC/sTG and the right retrosplenial hubs were significant for both effects simultaneously. In the other direction of the conjunction (for increased activity in the faces group) there were no voxels even at uncorrected levels.

These results do not contradict known findings about the effects of faces obtained in conventional analyses with the cognitive subtraction approach. In more posterior portions of the fusiform gyrus, more active during the task, the activation was higher in the faces group, as expected (10, −72, −2, *t* = 5.60, *p* = 0.005, peak-level corrected). Likewise, the amygdala was more active in the faces than in the mourning group (26, 2, −26, *t* = 4.66, *p* = 0.09). However, as the previous conjunction analysis has clarified, none of these relative activations were located within the faces evidence network.

### Effects of repetition of stimuli (repetition suppression) and its interaction with evidence

In the final analyses of this study, we considered the change of the signal over time. The task signal has been shown to decrease over the experiment as an effect of practice, and sensory and semantic association areas show signal decrements consistent with their recruitment during encoding (sensory priming, Schacter and Buckner 1998; see also Grill-Spector et al. 2006; Barron et al. 2016). Figure 8a simultaneously shows the evidence for faces network, the pattern of task activations and deactivations of Figure 5, and the decrements of this task activation elicited by regressing the data on the number of times the items of the trial had been previously displayed (green hatch pattern; Table A4). In the posterior parietal/occipital region and in the fusiform gyrus, decrements consistent with a priming effect were visible in the activated areas (Henson et al. 2002), adjacent to and partially overlapping the inferior parietal hub of the evidence for faces network. Decrements over time were also observed in activated areas in the prefrontal cortex and in the ventral striatum/brainstem. These latter are known effect respectively of practice or conceptual priming (Schacter and Buckner 1998) and novelty (Bunzeck and Düzel 2006) and will not be commented further here.

When looking at the interaction between the effect of evidence and repetition of stimuli, we found a third order interaction evidence × group × repetition (Table A5), shown in Figure 8a and b as a violet hatch pattern. In the parietal region, this interaction can be seen to involve the deactivated areas adjacent to but distinct from those of the evidence for the faces hubs (PGa and PFm, Geyer et al. 1999). In the retrosplenial hub, the interaction appeared as a distinct locus in the posterior cingulus (Figure 8b), which however failed to reach significance after correction. The most extensive areas affected by this third order interaction were in the medial aspect of the brain, where it modulated task deactivations that hosted hubs of the common decision network. The strongest effects here were in the vmPFC and dorsomedial prefrontal cortex (dmPFC). We had seen the posterior cingulus and vmPFC involved by connectivity patterns in Figure 4 (see panel a, connectivity at x = −8).

Separate inspection of the interaction evidence × repetition within the groups revealed the third order interaction to arise entirely from the mourning scenes task, which showed a much more complex encoding of the evidence for sadness. In the faces group, the effect of evidence was stable across the whole experiment, giving no interaction with the effect of repetition. In contrast to faces, in the mourning scenes task the hubs were encoding evidence for the decision only briefly at the beginning of the experiment (Figure 8c). In the last part of the experiment, encoding was characterized by activity for neutral scenes, which precluded any simple mapping of high and low evidence trials to activity levels.

## Discussion

In both groups, regressing the signal on the evidence for the decision revealed a network of areas centered in the somatosensory association cortex and the middle insula. In line with the alternative view, a cluster of the common decision network included the left precentral gyrus, which is consistent with the possibility that participants were accumulating evidence for button pressing (Cisek and Kalaska 2010). Deciding based on facial expressions, which offer rich sensory information for this task, additionally recruited the inferior parietal and postsplenial cortices, supporting the notion that recruitment of evidence hubs is to some extent selective. Furthermore, the selectivity of the face hubs was supported by connectivity data suggesting the existence of at least two partially segregated networks accumulating evidence for the decision. These findings are consistent with a generalized mechanism of evidence-accumulation, active also when deciding in an elementary social cognition task.

The hubs detected by modelling evidence for the decision are also consistent with data from cortical lesions. The main hub of the decision network recruited in both groups was located in the somatosensory association cortex, an area also implicated in studies of patients with impairments in classifying emotions (even if in lesions studies the effects are prevalently right-lateralized, Adolphs et al. 1996, 2000). The same studies also show a locus associated with impairment in the anterior fusiform gyrus (Adolphs et al. 1996), as in our data. The effect of lesions in the posterior iPC/sTG, the site of the faces evidence hubs and an area involved in the appraisal of emotional stimuli in imaging studies (Viviani 2013), are impairments in the computation of “a mental representation of extrapersonal events in terms of their motivational salience” (Mesulam 1999).

The goal of identifying decision hubs associated specifically with the mourning scenes, which required considering context, was more elusive. Deciding on scenes of mourning individuals activated specific evidence hubs, not recruited faces, but only briefly at the beginning of the task. These hubs included the inferior parietal cortex, an area often active in reappraisal of emotional stimuli (Viviani 2013; Messina et al. 2015) and extended or complemented the evidence hubs in both groups in the posterior cingulus and vmPFC, which have been assigned to a supramodal system based on their histological classification (Chanes and Barrett 2016). One possibility is that the fleeting character of evidence accumulation here was related to the contextual nature of the information in question.

The course of the signal showed that the lack of activation was due to a brief relative deactivation of variable intensity at the presentation of the stimuli in the evidence hubs. This deactivation then progressed to activation at the time of the decision in trials where the evidence of the decision was large. This pattern is visible in other decision making studies that report the signal course (Boorman et al. 2009; Viviani et al. 2017). In contrast, in the adjacent constantly task-deactivated areas, the deactivation remained stable. In a sensory-based decision making study, Tosoni et al. (2008) showed that stably task-deactivated areas, attributed to the default network system, were distinct from those modulated by decision making. Their findings are consistent to those obtained here in so far as the parietal hubs, while contributing to modulating the activation in the faces vs. mourning contrast, were adjacent to but did not completely engulf the task deactivations. The initially activated evidence hubs in the mourning group, however, were located within the deactivations of the default network. The relationship of areas such as vmPFC and the inferior parietal cortex, often active in tasks of social cognition and emotion processing, and the default network has been noted before (Mars et al. 2012; Viviani 2014).

As whole, the fMRI signal in posterior part of the brain appeared to be organized in large portions of cortex characterized by a gradient of progressive deactivation. Sensory areas, strongly activated by the task, were succeeded by associative areas modulated by sensory priming. The evidence hubs selectively recruited by decision about faces were located at the transition area to task deactivations. The progression ended in the deactivated areas of the default network system. Areas associated with differential task activation identified in previous studies that followed the cognitive subtraction approach were present also in our data. This was the case for the activation of intermediate portion of the fusiform gyrus (Kanwisher et al. 1997) and of the amygdala (Whalen et al. 1998) in the faces task. However, these activations did not completely characterize the pattern of the fMRI signal.

These results integrate the gradient model of cortical organization in the posterior and limbic portion of the brain (Huntenburg et al. 2018). Our data provide evidence that a gradient of task activation is associated with the progressive processing of information in a trajectory from sensory to supramodal association areas hosting the evidence hubs. They are also consistent with precursor models of brain function suggesting that high-level association areas are organized as sparse interconnected parallel systems (Goldman-Rakic 1988). Furthermore, they document the functional relationship between the areas at the top of the processing hierarchy (the evidence hubs) and task deactivations, substantiating the role of default network areas within this organizational gradient (Margulies et al. 2016).

If the organization of information processing in the posterior part of the brain has a neural correlate consisting of a gradient of progressive deactivation, questions arise about why this may be the case. While several related models have been developed in the psychological literature to explain empirical findings in decision making, it has been shown that all these models share a common fundamental computational step (Bogacz et al. 2006). Among the proposals for the implementation of this mechanism are several variations of competition between representations of the choice alternatives via mutual inhibition (Usher and McClelland 2001; Shadlen and Newsome 2001; Wang 2002; Cisek 2006). For this reason, evidence for decrements of the fMRI signal in preference-based choice tasks has been interpreted as evidence for this inhibition (Hunt et al. 2015), possibly mediated by GABA transmission (Jocham et al. 2012; for discussion, see Hunt and Hayden 2017). Since competition via mutual inhibition is thought to be a characteristic feature of neural circuits, one would expect decrements due to this mechanism to be present also in other choice tasks, as was the case here.

A similar conclusion may be reached when considering the theoretical literature on general models of information processing. According to the predictive coding theory of brain function, the cortex generates a predictive model of the sensory inputs and an inference about their causes (Rao and Ballard 1999; Friston 2005; Kilner et al. 2007; Friston et al. 2013). The areas higher in the hierarchy of processing feed predictions to lower areas. In this perspective, the decision hubs detected here are computing the decision evidence signal by matching the predicted distribution of the signal of interest in the external world (in the present case, the inferred degree of sadness of the displayed individuals). The same distribution in the pre-motor areas provides the match at the level of the response. According to this model, the sensory and associative areas would be tuned by feedback so as to provide the signal in support of this prediction (referred to as a ‘prediction error’ relative to the invariant signal common to all stimuli). As a result, only the information that is relevant for the target classification of input is present in the activity of the associative areas. These tuned cortical areas may correspond to those where neural priming is observed (Henson 2003; Auksztulewicz and Friston 2016), and that were here located between the early sensory processing areas and the evidence hubs.

In algorithmic implementations of predictive coding, where the relevant information in the input channel is represented by different pools of hidden units in a network, training is constrained by minimizing the amount of information in these units (‘minimal description length’, Hinton and Zemel 1994). This increases the redundancy of the codes represented by the activation states of decision nodes. This account draws attention to the fact that in the evidence hubs the amount of information collected from upstream areas may be minimal, encoding only a position between two possible points (the two alternative decisions, here indexed by the relative degree of sadness; cf. Koay et al. 2019). This makes the units that encode the relative degree of sadness to have similar activity, explaining why they may be all silent when the evidence for prediction is low. In contrast, early in the sensory processing channel the amount of information reflects relevant details of the visual input, requiring larger amounts of information to be coded. Also in this model, neural implementations are thought to make use of competitive inhibition to enforce progressive reduction in the amount of information (Hinton 2007). The minimal description length account is also consistent with the relatively low differentiation of cortical layers at the high end of the cortical gradient (Chanes and Barrett 2016; Huntenburg et al. 2018).

In these models, deactivation (increased inhibition) cannot be equated with lack of recruitment, as in the cognitive subtraction approach. This is exemplified by the present findings in the occipito-temporal cortex near the iPC/sTG hub and in the fusiform gyrus. In the posterior portion of the fusiform gyrus, where more elementary features of the visual input may be expected to be processed, task activity was larger in the faces group, which may be expected to make more intensive use of this type of visual information. Anteriorly, however, the evidence hub, where the evidence for the decision represents a high-level type of information, was located adjacent to the transition area to task deactivations and showed less task activity in the faces group, consistently with a stronger inhibitory influence.

If the progressive deactivation is related to inhibitory processes that constrain the complexity of representations of the encoded stimuli, the possibility arises that deactivated areas may host a parallel system that, operating side by side the activated centres of the executive dorsal network, organizes encoding of information. Whether these conjectures on the origin of deactivation are correct or not, much appears to take place in the weakly active or relatively deactivated ventral associative areas of the brain that is not captured by traditional analysis approaches and has received comparatively little attention as an imaging phenotype.

## Conclusion

In an elementary social cognition task a set of sparse areas spanning the parietal, temporal, and insular lobes were associated with the evidence for the decision, confirming the generality of the mechanism behind decision making. Moreover, these areas appeared to be organized in parallel networks that were differentially recruited depending on the information available for the decision and were embedded in gradients of increasing deactivations in associative areas. Notwithstanding the different starting point, these findings are consistent with morphological and computational accounts of brain function. Our data suggested that deactivations may be produced by the process of evidence accumulation, providing new hypotheses on the relationship between abstract cognition and the task deactivations of the default network system.

## Acknowledgments

This work was supported in part by a collaborative grant of the Federal Institute for Drugs and Medical Devices (BfArM, Bonn, Grant No. V-17568/68502/2017-2020) to Roberto Viviani. The authors declare no conflict of interest.

## Appendix

**Table A1.** Evidence.

| Cl. # | Brain area | MNI coord. | *k* | *p* (cl.) | *t* | *p* (peak) |
| --- | --- | --- | --- | --- | --- | --- |
| 1 | Left Inferior Parietal/Supramarginal (BA 40) | -56 -26 26 | 1055 | 0.007 | 6.16 | <0.001 |
|  | Left Postcentral Gyrus (BA 1) | -54 -20 56 |  |  | 4.95 | 0.027 |
|  | Left Inferior Parietal Lobule (BA 40) | -60 -26 48 |  |  | 3.98 | 0.262 |
|  | Left Postcentral/Somatosensory (BA 1/43) | -62 -20 38 |  |  | 3.90 | 0.300 |
|  | Left Postcentral Gyrus (BA 3) | -44 -24 46 |  |  | 3.71 | 0.418 |
|  | Left Inferior Parietal Lobule (BA 40) | -58 -40 46 |  |  | 3.59 | 0.490 |
|  | Left Postcentral Gyrus (BA 3) | -42 -30 62 |  |  | 3.27 | 0.688 |
| 2 | Left Insula (BA 13) | -38 0 14 | 258 | 0.044 | 5.33 | 0.007 |
|  | Left Primary Somatosensory Cortex (BA 43) | -54 -4 10 |  |  | 4.17 | 0.183 |
| 3 | Right Supramarginal Gyrus (BA 40) | 64 -36 42 | 466 | 0.019 | 4.98 | 0.024 |
|  | Right Supramarginal Gyrus (BA 40) | 66 -26 30 |  |  | 4.62 | 0.062 |
|  | Right Rolandic Operculum (BA 43)/Supra-marginal Gyrus (BA 48) | 52 -22 24 |  |  | 4.07 | 0.222 |
| 4 | Right Insula (BA 13) | 38 2 14 | 101 | 0.125 | 4.59 | 0.066 |
| 5 | Left Precentral Gyrus (BA 6) | -32 -18 66 | 89 | 0.140 | 4.21 | 0.166 |

**Legend.** MNI coordinates: Montreal Neurological Institute coordinates, in mm; *k*: cluster size, in 2x2x2 mm voxels; *p* (cl.): significance level, cluster-level correction; *t*: peak *t*; *p* (peak): significance level, peak-level correction. The table lists up to 8 peaks per cluster, at minimum distance 8mm.

**Table A2.** Evidence x faces.

| Cl. # | Brain area | MNI coord. | *k* | *p* (cl.) | *t* | *p* (peak) |
| --- | --- | --- | --- | --- | --- | --- |
| 1 | Left Parieto-occipital Gyrus (BA 39) | -42 -72 28 | 475 | 0.018 | 5.54 | 0.004 |
|  | Left Parieto-occipital Gyrus (BA 39) | -50 -72 22 |  |  | 5.19 | 0.012 |
| 2 | Right Precuneus (BA 23) | 16 -58 24 | 306 | 0.036 | 5.54 | 0.004 |
| 3 | Left Precuneus (BA 31) | -10 -62 24 | 306 | 0.036 | 4.59 | 0.070 |
|  | Left Retrosplenial Cortex (BA 29) | -8 -48 4 |  |  | 3.61 | 0.482 |
| 4 | Left Parahippocampal Gyrus (BA 27) | -30 -42 -8 | 93 | 0.137 | 4.24 | 0.164 |
|  | Left Fusiform Gyrus (BA 36/37) | -34 -36 -18 |  |  | 3.82 | 0.356 |
| 5 | Right Angular/Parieto-occipital (BA 39/19) | 48 -70 22 | 285 | 0.040 | 4.23 | 0.168 |
|  | Right Angular Gyrus (BA 39) | 42 -70 32 |  |  | 4.05 | 0.242 |
| 6 | Right Fusiform Gyrus (BA 37) | 34 -36 -16 | 42 | 0.252 | 3.92 | 0.308 |

**Legend.** MNI coordinates: Montreal Neurological Institute coordinates, in mm; *k*: cluster size, in 2x2x2 mm voxels; *p* (cl.): significance level, cluster-level correction; *t*: peak *t*; *p* (peak): significance level, peak-level correction. The table lists up to 8 peaks per cluster, at minimum distance 8mm.

**Table A3.** Conjunction Evidence x faces AND Task x faces (mourning vs. faces)

| Cl. # | Brain area | MNI coord. | *k* | *p* (cl.) | *t* | *p* (peak) |
| --- | --- | --- | --- | --- | --- | --- |
| 1 | Left Parieto-occipital Gyrus (BA 39) | -42 -68 24 | 455 | 0.036 | 4.25 | 0.343 |
| 2 | Right Precuneus (BA 23) | 18 -54 22 | 158 | 0.158 | 5.11 | 0.040 |
| 3 | Left Precuneus (BA 31) | -12 -60 22 | 233 | 0.097 | 4.38 | 0.266 |
| 4 | Left Parahippocampal Gyrus (BA 27) | -30 -42 -8 | 136 | 0.192 | 4.24 | 0.351 |
| 5 | Right Angular/Parieto-occipital (BA 39/19) | 44 -72 18 | 20 | 0.701 | 3.43 | 0.885 |
| 6 | Right Fusiform Gyrus (BA 37) | 34 -36 -16 | 60 | 0.408 | 3.92 | 0.582 |

**Legend.** MNI coordinates: Montreal Neurological Institute coordinates, in mm; *k*: conjunction cluster size, in 2x2x2 mm voxels; *p* (cl.): significance level, cluster-level correction; *t*: peak conjunction *t*; *p* (peak): significance level, peak-level correction. The cluster numbers correspond to those in Table A2.

**Table A4.** Repetition (decreases)

| Cl. # | Brain area | MNI coord. | *k* | *p* (cl.) | *t* | *p* (peak) |
| --- | --- | --- | --- | --- | --- | --- |
| 1 | Right Inferior Frontal Gyrus (BA 45) | 46 8 26 | 6197 | 0.001 | 8.72 | <0.001 |
|  | Right Anterior Insula (BA 44/47) | 32 24 0 |  |  | 7.68 | <0.001 |
|  | Right Nucleus Accumbens | 12 6 -2 |  |  | 7.25 | <0.001 |
|  | Left Premotor Cortex/Supplementary Motor Cortex (BA 6) | -6 20 46 |  |  | 7.00 | <0.001 |
|  | Right Inferior Frontal Gyrus (BA 45) | 50 28 16 |  |  | 6.84 | <0.001 |
|  | Right Supplementary Motor Cortex (BA 6) | 8 12 52 |  |  | 5.72 | 0.003 |
|  | Right Precentral Gyrus (BA 6) | 40 2 42 |  |  | 4.97 | 0.027 |
| 2 | Left Anterior Insula (BA 44/47) | -30 22 2 | 4835 | 0.001 | 8.38 | <0.001 |
|  | Left Premotor Cortex/Supplementary Motor Cortex (BA 6) | -48 10 26 |  |  | 6.95 | <0.001 |
|  | Left Inferior Frontal Gyrus (BA44/45) | -44 18 24 |  |  | 6.83 | <0.001 |
|  | Left Inferior Frontal Gyrus (BA 44) | -40 6 32 |  |  | 6.77 | <0.001 |
|  | Left Nucleus Accumbens | -10 6 -4 |  |  | 6.70 | <0.001 |
|  | Left Precentral Gyrus (BA 6) | -38 -2 56 |  |  | 6.28 | <0.001 |
|  | Left Inferior Frontal Gyrus (BA 45) | -46 24 14 |  |  | 5.82 | 0.002 |
|  | Left Subgenual Ventromedial Prefrontal Cortex (BA 25) | -14 0 -10 |  |  | 5.54 | 0.005 |
| 3 | Right Lingual Gyrus (BA 19) | 24 -60 -6 | 3556 | 0.001 | 7.71 | <0.001 |
|  | Right Middle Temporal Gyrus (BA 21) | 48 -62 12 |  |  | 6.00 | 0.001 |
|  | Right Fusiform Gyrus (BA 37) | 30 -38 -14 |  |  | 5.57 | 0.004 |
|  | Right Fusiform Gyrus (BA 37) | 42 -46 -16 |  |  | 5.38 | 0.009 |
|  | Right Inferior/Middle Temporal Gyrus (BA 20/21) | 48 -64 -2 |  |  | 5.37 | 0.009 |
|  | Right Superior Occipital Gyrus (BA 19) | 26 -74 36 |  |  | 5.29 | 0.011 |
|  | Right Lingual Gyrus (BA 19) | 24 -46 -8 |  |  | 5.12 | 0.017 |
|  | Right Middle Occipital Gyrus (BA 19) | 36 -82 24 |  |  | 4.87 | 0.034 |
| 4 | Left Fusiform Gyrus (BA37) | -26 -44 -10 | 4469 | 0.001 | 7.14 | <0.001 |
|  | Left Fusiform Gyrus (BA37) | -28 -56 -10 |  |  | 7.08 | <0.001 |
|  | Left Fusiform Gyrus (BA37) | -32 -72 -8 |  |  | 5.99 | 0.001 |
|  | Left Middle Temporal Gyrus (BA21) | -48 -66 4 |  |  | 5.87 | 0.002 |
|  | Left Middle Occipital Gyrus (BA 19) | -24 -74 32 |  |  | 5.71 | 0.003 |
|  | Left Middle Occipital Gyrus (BA 19) | -30 -84 30 |  |  | 5.53 | 0.005 |
|  | Left Middle Temporal Gyrus (BA 39) | -44 -66 16 |  |  | 5.42 | 0.008 |
|  | Left Fusiform Gyrus (BA 37) | -42 -48 -18 |  |  | 5.07 | 0.020 |
| 5 | Left Middle Temporal Gyrus (BA 21) | -54 -6 -14 | 106 | 0.117 | 4.64 | 0.062 |
| 6 | Right Postcentral Gyrus (BA 3) | 44 -26 42 | 234 | 0.048 | 4.30 | 0.142 |
|  | Right Postcentral Gyrus (BA 3) | 56 -20 40 |  |  | 3.73 | 0.405 |
|  | Right Postcentral Gyrus (BA 3) | 36 -30 48 |  |  | 3.70 | 0.420 |
|  | Right Postcentral Gyrus (BA3) | 64 -22 34 |  |  | 3.64 | 0.455 |
| 7 | Left Postcentral Gyrus (BA 3) | -46 -26 38 | 176 | 0.070 | 4.20 | 0.180 |
|  | Left Supramarginal Gyrus (BA 40) | -32 -34 40 |  |  | 4.09 | 0.224 |
| 8 | Right Primary Somatosensory Cortex (BA7) | 26 -48 54 | 53 | 0.209 | 3.85 | 0.331 |

**Legend. MNI coordinates: Montreal Neurological Institute coordinates, in mm; *k*: cluster size, in 2x2x2 mm voxels; *p* (cl.): significance level, cluster-level correction; *t*: peak *t*; *p* (peak): significance level, peak-level correction. The table lists up to 8 peaks per cluster, at minimum distance 8mm.**

**Table A5.** Interaction evidence × repetition (decreases) × mourning.

| Cl. # | Brain area | MNI coord. | *k* | *p* (cl.) | *t* | *p* (peak) |
| --- | --- | --- | --- | --- | --- | --- |
| 1 | Left Superior Temporal Gyrus (BA 22)/Angular Gyrus (BA 39) | -54 -54 20 | 1380 | 0.005 | 6.12 | <0.001 |
|  | Left Supramarginal Gyrus (BA 40) | -52 -58 30 |  |  | 5.87 | <0.001 |
|  | Left Inferior Parietal Lobule/Supramarginal Gyrus (BA 40) | -48 -50 40 |  |  | 4.75 | 0.041 |
| 2 | Left Middle Cingulate Cortex (BA 8) | -2 44 46 | 1587 | 0.004 | 5.61 | 0.003 |
|  | Left Middle Cingulate Cortex (BA 8) | -10 36 38 |  |  | 5.12 | 0.015 |
|  | Left Anterior Cingulate Cortex (BA 24) | -4 48 16 |  |  | 4.68 | 0.051 |
|  | Left Dorsomedial Prefrontal Cortex/Frontal Eye Field (BA 8) | -14 26 52 |  |  | 4.46 | 0.089 |
|  | Right Dorsolateral Prefrontal (BA 9) | 10 48 32 |  |  | 4.37 | 0.111 |
|  | Left Middle Frontal Gyrus (BA 46) | -10 36 50 |  |  | 4.22 | 0.159 |
|  | Left Middle Frontal Gyrus (BA 46) | -18 58 24 |  |  | 3.93 | 0.272 |
|  | Left Middle Frontal Gyrus (BA 46) | -4 54 30 |  |  | 3.88 | 0.294 |
| 3 | Right Supramarginal Gyrus (BA 40) | -38 18 44 | 296 | 0.038 | 4.55 | 0.069 |
|  | Right Supramarginal Gyrus (BA 40) | -48 18 36 |  |  | 3.72 | 0.383 |
| 4 | Left Anterior Cingulate (BA 24) | -46 48 -4 | 149 | 0.072 | 4.27 | 0.141 |
| 5 | Right Inferior Frontal Gyrus (BA 45) | 54 -56 32 | 355 | 0.031 | 4.13 | 0.188 |
|  | Right Superior Temporal Gyrus (BA 22)/Angular Gyrus (BA 39) | 58 -64 30 |  |  | 3.72 | 0.387 |
| 6 | Left Supramarginal Gyrus (BA 40) | -6 36 -4 | 75 | 0.145 | 4.03 | 0.224 |
| 7 | Left Inferior Parietal Lobule/Supramarginal Gyrus (BA 40) | -50 18 20 | 51 | 0.200 | 3.71 | 0.389 |

**Legend.** MNI coordinates: Montreal Neurological Institute coordinates, in mm; *k*: cluster size, in 2x2x2 mm voxels; *p* (cl.): significance level, cluster-level correction; *t*: peak *t*; *p* (peak): significance level, peak-level correction. The table lists up to 8 peaks per cluster, at minimum distance 8mm.

